# Neuronal mitochondrial morphology is significantly affected by both fixative and oxygen level during perfusion

**DOI:** 10.1101/2022.09.08.507147

**Authors:** Su Yeon Kim, Klaudia Strucinska, Bertha Osei, Kihoon Han, Seok-Kyu Kwon, Tommy L. Lewis

**Author notes:** These authors contributed equally to this work (Su Yeon Kim, Klaudia Strucinska, Bertha Osei).

## Abstract

Neurons in the brain have a uniquely polarized structure consisting of multiple dendrites and a single axon generated from a cell body. Interestingly, intracellular mitochondria also show strikingly polarized morphologies along the dendrites and axons: in cortical pyramidal neurons (PNs) dendritic mitochondria have a long and tubular shape, while axonal mitochondria are small and circular. Mitochondria play important roles in each compartment of the neuron by generating ATP and buffering calcium, thereby affecting synaptic transmission and neuronal development. In addition, mitochondrial shape, and thereby function, is dynamically altered by environmental stresses such as oxidative stress, or in various neurodegenerative diseases including Alzheimer’s disease and Parkinson’s disease. Although the importance of altered mitochondrial shape has been claimed by multiple studies, methods for studying this stress-sensitive organelle have not been standardized. Here we address the pertinent steps that influence mitochondrial morphology during experimental processes. We demonstrate that fixative solutions containing only paraformaldehyde (PFA), or that introduce hypoxic conditions during the procedure induce dramatic fragmentation of mitochondria both in vitro and in vivo. This disruption was not observed following the use of glutaraldehyde addition or oxygen supplementation, respectively. Finally, using pre-formed fibril α-synuclein treated neurons, we show a difference between mitochondrial morphology when samples were fixed with PFA/glutaraldehyde or PFA/sucrose containing solutions, but not PFA alone. Our study provides optimized methods for examining mitochondrial morphology in neurons, and demonstrates that fixation conditions are critical when investigating the underlying cellular mechanisms involving mitochondria in physiological and neurodegenerative disease models.

## INTRODUCTION

Neurons are unique cells with distinct sizes, highly complex morphologies and the ability to transmit and receive electro-chemical messages. The process of neurotransmission requires vast amounts of energy in the form of ATP (Rolfe and Brown 1997). While the respective contributions of the various metabolic processes to supply this ATP are not fully understood in neurons, mitochondria play a vital roles in the ability of neurons to properly regulate their energy needs, as well as other critical processes including Ca^2+^ handling, lipid biogenesis and regulation of the apoptotic pathway (Fox, Raichle et al. 1988, Belanger, Allaman et al. 2011, Chandel 2014, Diaz-Garcia and Yellen 2019).

The recent explosion in techniques to label and visualize organelles with high spatial resolution has revealed that excitatory PNs appear to contain distinct mitochondrial subpopulations within their respective cellular compartments of the axon, soma and dendrites. Axonal mitochondria are small, individual entities while dendrites contain mitochondria that are highly elongated and overlapping each other (Popov, Medvedev et al. 2005, Chang and Reynolds 2006, Lewis, Kwon et al. 2018, Rangaraju, Lauterbach et al. 2019). Mitochondria within the soma display intermediate size and morphology between those found in the axons and dendrites (Faitg, Lacefield et al. 2021, Turner, Macrina et al. 2022).

Although the majority of previous data meets this consensus, there are discrepancies in the range of sizes. We reported that in mouse layer 2/3 cortical PNs dendritic mitochondrial length ranges from 1.31 to 13.28μm long, but only 0.45 to 1.13 μm in axons (Lewis, Kwon et al. 2018). However, other studies observed both shorter dendritic mitochondria in cortical neurons (Kimura and Murakami 2014), or longer dendritic mitochondria in live imaged cultured hippocampal neurons (Rangaraju, Lauterbach et al. 2019). In addition, a recent study presented the length of dendritic mitochondria as 4.7 ± 0.9 μm by 3D electron microscopy (EM) imaging, but Ca^2+^ transients in the mitochondrial matrix extended to 15 μm (Lin, Li et al. 2019). The reason for these gaps in size have not yet been scrutinized.

Mitochondria are proposed to be critical sites of reduced efficiency and function during the processes of normal aging and pathogenic neurodegeneration (Swerdlow, Burns et al. 2014, Azzu and Valencak 2017). A common observation across many different forms of neurodegenerative diseases (including Alzheimer’s and Parkinson’s diseases) suggests significant reduction in mitochondria number and size as well as a loss of mitochondrial ultrastructure (Zhang, Trushin et al. 2016, Gonzalez-Rodriguez, Zampese et al. 2021). In addition, fragmented mitochondria are increased following cerebral ischemia (Owens, Park et al. 2015, Zhou, Chen et al. 2021).

Based on our own observations, and the large variance in the reported sizes of dendritic mitochondria in excitatory pyramidal neurons and fixation conditions in the literature, we hypothesized that standard fixation conditions may not faithfully capture the mitochondrial structure present in living neurons. Thus, we rigorously tested the effects of multiple fixation parameters on the morphology of mitochondria in both cultured primary PNs as well as PNs in vivo. We find that a combination of direct fixation with a mixture of paraformaldehyde (2%) and glutaraldehyde (0.075%) with the presence of sufficient oxygen are critical for maintaining mitochondrial morphology during fixation *in vitro* and *in vivo*.

## METHODS

### Animals

Animals were handled according to Institutional Animal Care and Use Committee (IACUC) approved protocols at the Oklahoma Medical Research Foundation (OMRF) and Korea Institute of Science and Technology (KIST-2020-133). Time-pregnant females of CD-1 IGS strain (Strain Code: 022) were purchased at Charles River Laboratories or Daehan Biolink (Eumseong, Korea) and used for in utero electroporation experiments and primary neuronal cultures.

### Plasmids

pCAG::mtYFP-P2A-tdTomato and pCAG::mt-YFP were previously published in (Lewis, Turi et al. 2016). pCAG::Cre and pCAG::HA-mCherry were previously used in (Kwon, Sando et al. 2016). pAAV EF1α::Flex Venus-T2A-mito-mScarlet was created by replacing Synaptophysin-Venus to Venus-T2A-mito-mScarlet in pAAV EF1α::Flex-Synaptophysin-Venus from (Kwon, Sando et al. 2016).

### Cell Lines

Mouse Embryonic Fibroblast (NIH/3T3) were purchased from ATCC (CRL-1658). 1×10^5^ cells of NIH/3T3 cells suspended in media (DMEM, gibco) with penicillin/streptomycin (0.5×; gibco) and FBS (sigma) were seeded on coverslips (Collagen Type I, Corning) in 6 well dishes. Transfection with plasmid DNA (1 mg/mL) using jetPRIME® reagent according to manufacturer protocol was performed 24 hours after seeding. Half of the coverslips were fixed with 2% PFA/0.075% GA in 1× PBS and the other half with 4% PFA in PBS 24 hours after transfection for 7 minutes. Each well with coverslip was washed three times with 1× PBS (sigma) for 10 minutes and mounted on microscope slides with Aqua PolyMount (PolyMount Sicences, Inc) and kept at 4 °C after drying overnight.

### Ex utero electroporation

A mix of endotoxin-free plasmid preparation (2 mg/mL) and 0.5% Fast Green (Sigma) mixture was injected using FemtoJet 4i (Eppendorf) into the lateral ventricles of isolated heads of E15.5 mouse embryos. Embryonic neural progenitor cells were electroporated using an electroporator (ECM 830, BTX) and gold paddles with four pulses of 20 V for 50 ms with 500 ms interval and an electrode gap of 1.0 mm. Dissociated primary neuron culture was performed after ex utero electroporation.

### Primary neuronal culture

Following ex utero electroporation, embryonic mouse cortices (E15.5) were dissected in Hank’s Balanced Salt Solution (HBSS) supplemented with HEPES (10 mM, pH 7.4), and incubated in HBSS containing papain (Worthington; 14 U/mL) and DNase I (100 μg/mL) for 15 min at 37 °C with a gentle flick between incubation. Samples were washed with HBSS three times, and dissociated by pipetting on the fourth wash. Cells were counted using Countess™ (Invitrogen) and cell suspension was plated on poly-D-lysine (1 mg/mL, Sigma)-coated glass bottom dishes (MatTek) or poly-D-lysine/laminin coated coverslips (BD bioscience) in Neurobasal media (Gibco) containing FBS (2.5%) (Sigma), B27 (1 X) (Gibco), and Glutamax (1 X) (Gibco). After 7 days, media was changed with supplemented Neurobasal media without FBS.

### Fixation for primary neuron culture

Half of the culture dishes were fixed with 2% paraformaldehyde (PFA) (PFA Alfa Aesar)/ 0.075% glutaraldehyde (GA) (Electron Microscopy Science, EMS) in 1× PBS (Sigma) and the other half was fixed with 4% PFA for 7 minutes. Dishes were washed three times with 1× PBS (sigma) for 10 minutes.

### In utero electroporation

A mix of endotoxin-free plasmid preparation (0.5 mg/mL) and 0.5% Fast Green (Sigma) was injected into one lateral hemisphere of E15.5 embryos using FemtoJet 4i (Eppendorf). Embryonic neural progenitor cells were labelled using the electroporator (ECM 830, BTX) with gold paddles at E15.5. Electroporation was performed by placing the anode (positively charged electrode) on the side of DNA injection and the cathode on the other side of the head. Five pulses of 38 V for 50 ms with 500 ms interval and an electrode gap of 1.0mm were used for electroporation.

### Intracardial perfusion

For direct and indirect perfusion experiments in Figure 2, animals were put to sleep using 5% isoflurane mixed with air and exsanguinated 21 days after birth (P21) by terminal intracardial perfusion. Pups were randomly divided in 4 groups, to test different perfusion conditions. 1x PBS and fixatives were kept on ice during the entire procedure. Group 1 (indirect PFA/GA) was perfused with 10 mL of 1x PBS followed by 30 mL of fixative 2% PFA/0.075% GA in PBS (32% PFA Alfa Aesar, 3% GA Electron Microscopy Science). Group 2 (indirect PFA) with 10 mL of PBS followed by 30 mL of 4% PFA in PBS. Group 3 (direct PFA/GA) by 30 mL of fixative 2% PFA/0.075% GA in PBS. Group 4 (direct PFA) 30 mL of 4% PFA in PBS. Animals were then dissected to isolate brains, that were later subjected to 20h post fixation in the same fixative that was used for perfusion in each group.

**Figure 1:**
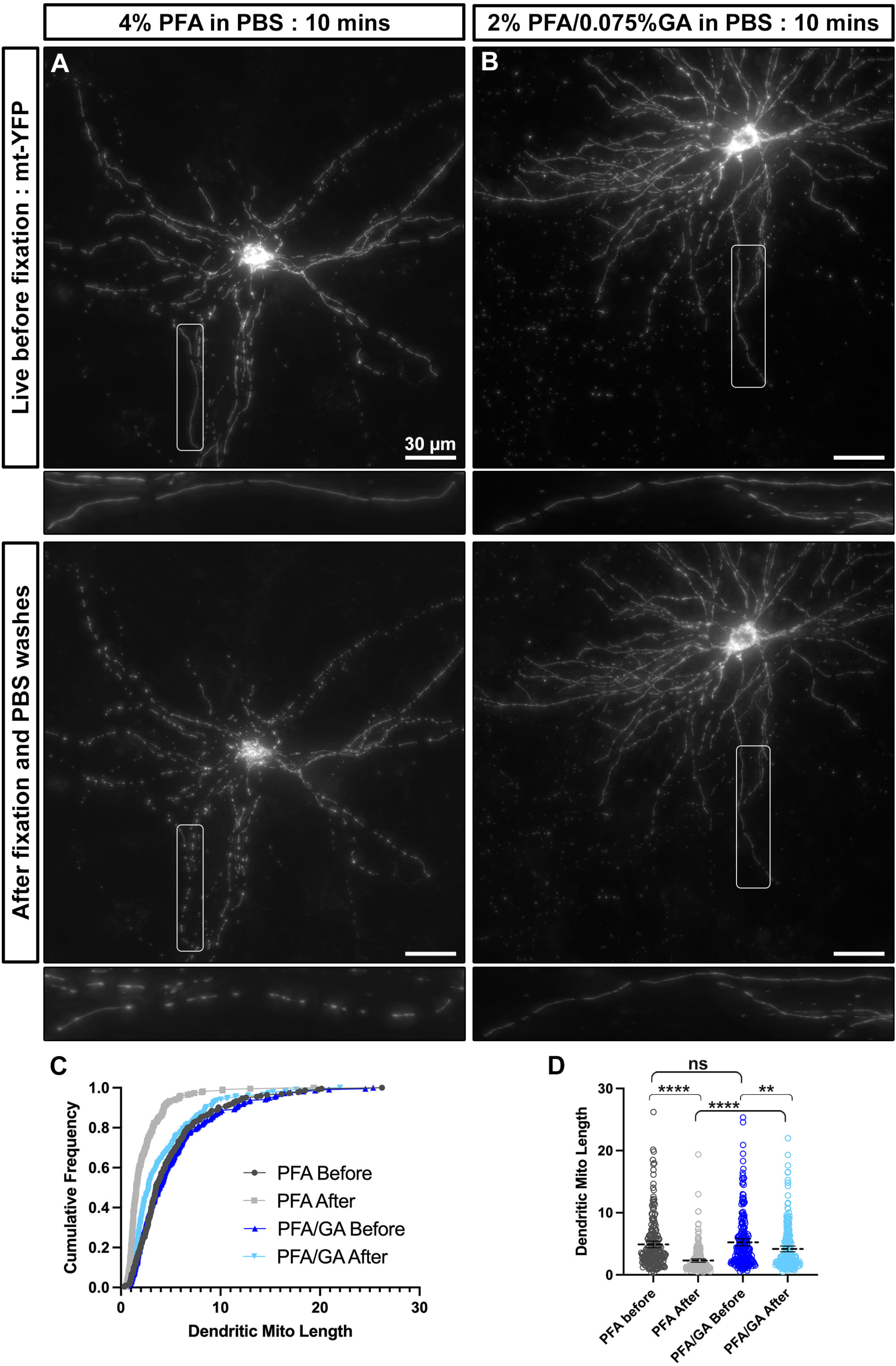
A combination of paraformaldehyde and glutaraldehyde maintains mitochondria morphology better than paraformaldehyde alone in cultured cortical neurons. A) A representative neuron labeled with mitochondrial matrix targeted YFP (mt-YFP) imaged live before fixation (top) and after fixation with 4% PFA in PBS (bottom). B) A representative neuron labeled with mt-YFP imaged live before fixation (top) and after fixation with 2% PFA/0.075% GA in PBS (bottom). C) Cumulative frequencies of dendritic mitochondrial length showing that fixation with PFA only results in fragmentation of mitochondria. D) Quantification of dendritic mitochondrial lengths shown as Mean +/− 95% CI before and after fixation with the indicated fixative solution. Kruskal–Wallis test with Dunn’s multiple comparisons test. n_4%PFAbefore_= 2 cultures, 16 dendrites, 225 mitochondria, n_4%PFAafter_= 2 cultures, 16 dendrites, 227 mitochondria, n_PFA/GAbefore_= 2 cultures, 16 dendrites, 222 mitochondria, n_PFA/GAafter_= 2 cultures, 16 dendrites, 238 mitochondria. **p<0.01, p*** p<0.001, **** p<0.0001. Scale bar = 30μm

**Figure 2:**
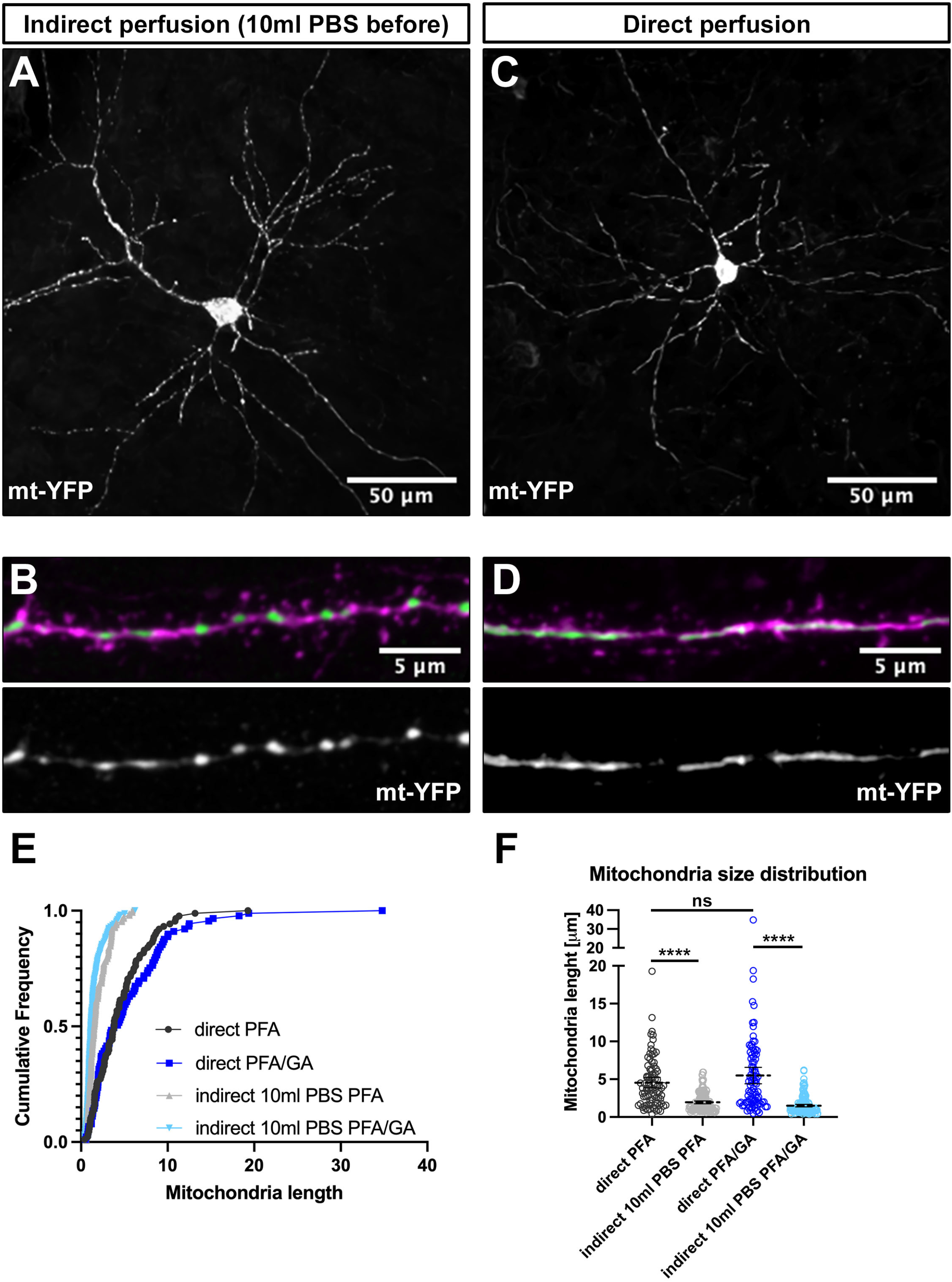
Direct perfusion is necessary for preserving mitochondrial morphology in vivo. A) Low magnification representative neuron following indirect perfusion with PFA/GA solution. B) High magnification of a dendrite segment following indirect perfusion with PFA/GA. C) Low magnification representative neuron following direct perfusion with PFA/GA solution. B) High magnification of a dendrite segment following direct perfusion with PFA/GA. C) Cumulative frequencies of dendritic mitochondrial length showing that indirect perfusion results in fragmentation of mitochondria. D) Quantification of dendritic mitochondrial lengths shown as Mean +/− 95% CI before and after fixation with the indicated fixative solution. Kruskal–Wallis test with Dunn’s multiple comparisons test. n_indirectPFA_= 15 dendrites, 150 mitochondria, n_directPFA_= 11 dendrites, 88 mitochondria, n_indirectPFA/GA_ = 13 dendrites, 158 mitochondria, n_directPFA/GA_ = 14 dendrites, 88 mitochondria. ns – not significant, **** p<0.0001. Scale bar = 50μm for A and C, 5μm for B and D

For oxygen supplement related experiments in Figure 3, mice were randomly assigned to anesthetize either with 2.5 vol% isoflurane with 2 ml/min oxygen or with isoflurane only. For the group with oxygen inhalation, isoflurane was delivered with oxygen through an anesthesia machine. A closed jar containing 1ml of isoflurane was used to anesthetize the group without oxygen. Mice were perfused transcardially with 2% PFA (Alfa Aesar) and 0.075% GA (Sigma) in PBS and brains were isolated for further experiments. During the perfusion, pulse and oxygen concentration were measured with MouseSTAT® Jr.Pulse Oximeter & Heart Rate Monitor (Kent Scientific). After 2 hours of post fixation in the same fixative used in perfusion, brains were washed with PBS and sectioned using a vibratome (Leica VT1200) at 130 μm. Sections were then washed 3 times with PBS and mounted on slides with VECTASHIELD® Vibrance™ Antifade Mounting Medium with DAPI (Vector laboratories).

**Figure 3:**
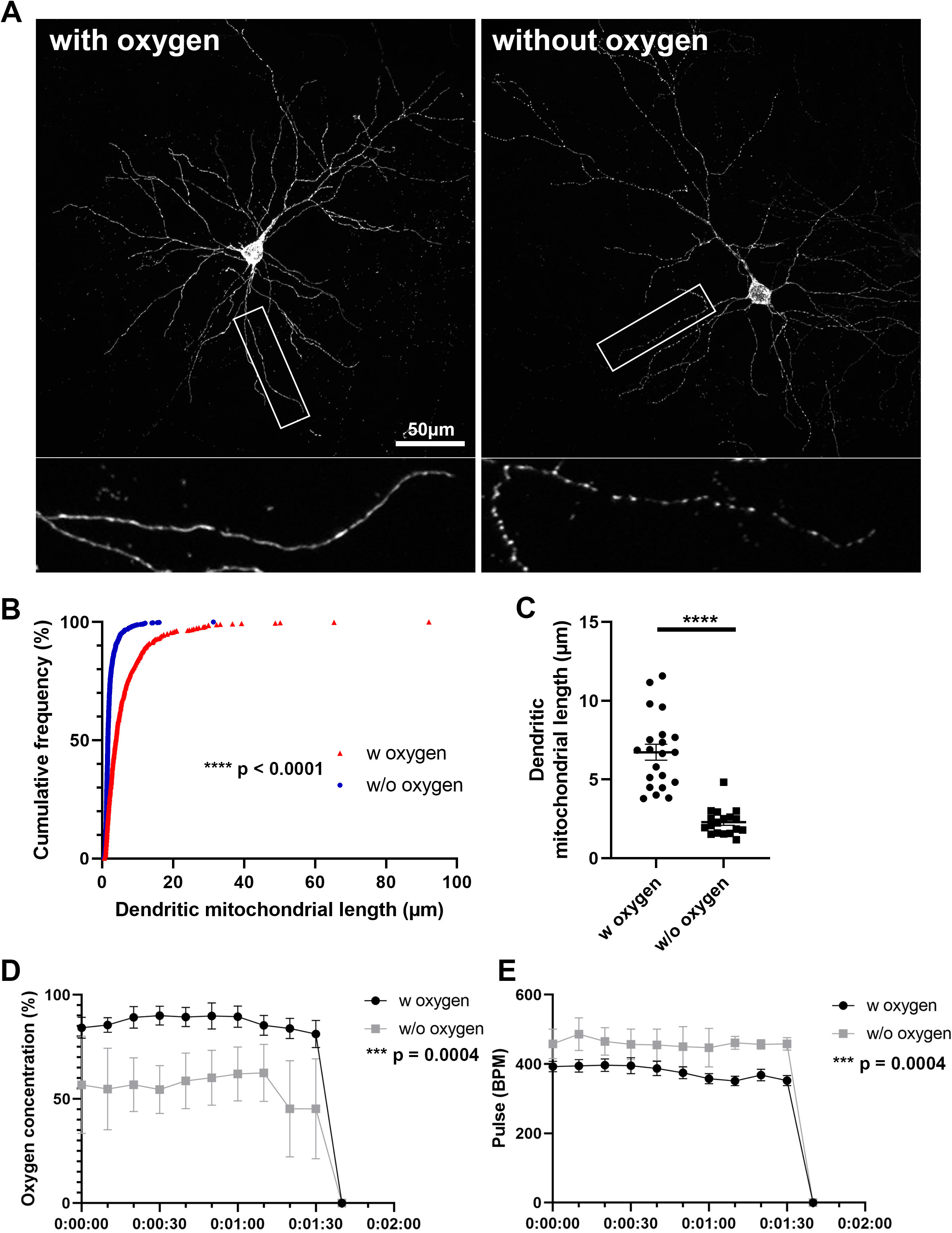
Oxygen supply is necessary for preserving mitochondrial morphology during anesthesia and perfusion. A) Representative images of mitochondria after perfusion. B) Cumulative frequencies of dendritic mitochondrial length showing that perfusion without oxygen supplement results in fragmentation of mitochondria. n_with oxygen_ = 627 mitochondria, n_without oxygen_ = 1061 mitochondria. Kolmogorov–Smirnov test, **** p<0.001 C) Quantification of dendritic mitochondria in each perfusion condition confirmed shorter mitochondria in perfusion without oxygen compared to perfusion with oxygen. n_with oxygen_ = 42 dendrites, n_without oxygen_ = 36 dendrites. unpaired t-test, **** p<0.0001 D & E) Oxygen concentration and pulse were measured during perfusion procedure. N_with oxygen_ = 7 mice, N_without oxygen_ = 6 mice. Mann-Whitney test, *** p=0.0004. Scale bar = 50μm.

### Imaging of cultured NIH3T3 cells and cultured neurons

Cultured cells were imaged on a Nikon Ti2 widefield system equipped with a Hammamatsu Fusion CMOS camera, standard cubes for FITC, TRITC and DAPI, and 60× (1.2NA) oil objective, and live imaging chamber from OXO. The whole system is controlled by Nikon Elements. For cultured NIH3T3 cells, cells were fixed as above and then imaged directly after fixation. For cultured neurons, cells were imaged live first and then fixed as above and then the same cell reimaged following fixation.

### Live imaging under hypoxic conditions

Live cell imaging under hypoxic conditions was performed with the EVOS M7000 imaging system (Thermo Scientific) equipped with an onstage incubator. Normal tyrode solution was used as a bath solution and neurons were incubated in a humidified atmosphere containing 5% CO_2_ at 37°C. To induce the hypoxic condition, oxygen level was gradually decreased from 20% to 10%, 5% and 0.1%. Samples were imaged for 10min with 1 min interval in each oxygen level. 1 image per oxygen level was used to analyze the dendritic mitochondrial length in each condition.

### Imaging brain sections

For direct and indirect perfusion experiments in Figure 2, fixed samples were imaged on a Zeiss LSM 880 confocal microscope controlled by Zeiss Black software. Imaging required two lasers 488nm and 561nm together with Zeiss objectives 40× (1.2NA) with 2× zoom, or 100× oil (1.25NA) with 3× zoom.

For oxygen supplement related experiments in Figure 3, fixed samples were imaged on a Nikon A1R confocal microscope with a Nikon objective 60× (1.25NA). Samples were visualized by Z-stacking that was later processed into Maximum Intensity Projection (MIP). MIP 2D images were then used for analysis of mitochondrial length and occupancy using NIS Elements software (Nikon) and Fiji (Image J).

### α-Synuclein treatment

Active human recombinant α-synuclein preformed fibrils (PFF) were purchased from StreeMarq. Before the treatment, α-synuclein PFF was diluted in PBS at 0.1mg/ml and sonicated for 30 sec. α-Synuclein was added at 10 DIV in a concentration of 1μg/ml and incubated for 11 days.

### Immunohistochemistry

Brains were washed 3 times for 15 minutes in 1× PBS and embedded in 3% low melt agarose (RPI, A20070) in 1× PBS. Brains in agarose cubes were sectioned using a vibratome (Leica VT1200) at 120 μm. Sections were then incubated with primary antibodies (chicken anti-GFP Aves Lab 1:1000, rabbit anti-dsRed Abcam 1:1000) that were diluted in the Blocking buffer (1%BSA, 0.2%TritonX-100, 5%NGS in PBS) at 4 °C for 48h. Subsequently sections were washed 6 times for 10 min in PBS and incubated with secondary antibodies (Alexa conjugated goat anti-chicken488 and goat anti-rabbit568 1:1000) at 4 °C for 48h. The excess of secondary antibodies was removed by six 10 minutes washes in 1× PBS. In the end sections were mounted on slides and coversliped with Aqua PolyMount (PolyMount Sicences, Inc.) and kept at 4 °C.

### Immunocytochemistry

Cultured neurons were fixed for 15 min at room temperature in either 4% PFA, 2% PFA with 0.075% GA, or 4% PFA with 4% sucrose and then washed with PBS. Cells were permeabilized with 0.2% Triton X-100 in PBS and incubation in 0.1% BSA and 2.5% goat serum in PBS was followed to block nonspecific signals. Primary and secondary antibodies were diluted in the blocking buffer described above and incubated at 4°C overnight. Coverslips were mounted on slides with VECTASHIELD® Vibrance™ Antifade Mounting Medium (Vector laboratories). Primary antibodies used in this experiment were mouse anti-HA (Biolegend, 1:500) and rabbit anti-α-Synuclein (pS129) (Abcam, 1:300), and all secondary antibodies were Alexa-conjugated (Invitrogen) and used at 1:1000 dilution.

### Quantification and statistical analysis

Statistical analysis was done in GraphPad’s Prism 6. Statistical tests, p-values, and (n) numbers are presented in the figure legends. Gaussian distribution was tested using D’Agostino & Pearson’s omnibus normality test. We applied non-parametric tests when data from groups tested deviated significantly from normality. No blinding was performed. No sample size calculation was performed. No exclusion critera were pre-determined and no animals were excluded. All analyses were performed on raw imaging data without any adjustments. Images in figures have been adjusted for brightness and contrast (identical for control and experimental conditions in groups compared), and images in Figure 2 have been processed with Nikon’s proprietary denoise.ai for visualization purposes only.

## RESULTS

### Fixation of cultured cells with a solution of PFA/GA better preserves mitochondrial morphology

Based on our own observations and the high variation that has been reported in the literature for mitochondrial morphology in neurons, we hypothesized that standard fixation conditions with 4% paraformaldehyde (PFA) may not be optimal for maintaining mitochondrial structure. To test this hypothesis, we performed ex utero electroporation (EUE) coupled with primary neuron culture to visualize mitochondrial morphology at single cell resolution in culture. Following culture for 17 days to allow for neuronal maturation, we first imaged neurons live at 37°C (**Figure 1A** (top) followed by immediate fixation with a 4%PFA solution in PBS and three PBS washes on the stage allowing us to image the same neurons following fixation (**Figure 1A** (bottom). As published previously by multiple groups, dendritic mitochondria are highly elongated and tubular in cortical PNs (Popov, Medvedev et al. 2005, Chang and Reynolds 2006, Lewis, Kwon et al. 2018, Rangaraju, Lauterbach et al. 2019) and occupy a large portion of the dendritic arbor in living neurons. However, upon fixation with 4% PFA, we observed a rapid fragmentation of dendritic mitochondria leading to smaller mitochondria (live dendritic mitochondria mean length: 4.9 ± 0.27μm; after 4% PFA fixation: 2.3 ± 0.14μm). This result confirmed that fixation with only a solution of 4% PFA is not optimal for maintaining the mitochondrial morphology observed in living neurons. In an attempt to determine a perfusion solution that would better preserve mitochondrial morphology during fixation, we searched the literature and observed that many groups performing electron microscopy included glutaraldehyde (GA) in their fixation solution as it provides increased cross-linking activity compared to PFA alone. To test if a solution of PFA/GA would achieve more optimal preservation of mitochondrial morphology, we performed the same experiment as above but used a fixative solution of 2%PFA/0.075%GA to fix the cultured neurons (**Figure 1B**). As clearly observed and quantified (**Figure 1C-D**), a solution of 2%PFA/0.075%GA dramatically reduced mitochondrial fragmentation during fixation (live dendritic mitochondria mean length: 5.2 ± 0.29μm; after PFA/GA: 4.2 ± 0.23μm). Finally, we asked if this was a result specific to the fixation of neuronal mitochondria or if it would be conserved in other cell types. Using cultured NIH3T3 cells transfected to fluorescently labeled mitochondria, we compared fixation with either 4%PFA or a solution of 2%PFA/0.075%GA (**Figure S1**). We observed very similar results to neurons with 4%PFA leading to a decrease in total mitochondrial area (4%PFA: 105.3μm^2^ vs 188.4μm^2^ for 2%PFA/0.075%GA) and increased circularity (4%PFA: .78 ± 0.01 vs .58 ± 0.01 for 2%PFA/0.075%GA).

### Direct perfusion results in the consistent preservation of mitochondrial morphology

Following this observation in cultured neurons, we tested if it applies to preservation of mitochondrial morphology in neurons *in vivo*. To test this, we performed *in utero* electroporation (IUE) on E15.5 CD1 pups with a plasmid encoding a mitochondrial matrix-targeted yellow fluorescent protein (mt-YFP). At postnatal day 21, when several aspects of neuronal differentiation, including dendritic morphogenesis and synaptogenesis, are adult-like, mice were anesthetized with isoflurane and intracardiac perfusion performed. Initial attempts resulted in a significant degree of mitochondrial fragmentation or loss of elongated mitochondria in perfused brains regardless of fixation solution (**Figure 2A-B,E-F**). We then tested various buffers, buffer pH and buffer temperature but none preserved the elongated mitochondria observed in the dendrites (data not shown). Finally, we attempted direct perfusion where instead of performing a pre-flush with PBS or other buffer to remove blood we directly started the perfusion with the fixative solution. Strikingly, this resulted in consistent preservation of the elongated mitochondria morphology observed in living neuronal dendrites (**Figure 2C-D, E-F**). While PFA/GA still resulted in the most elongated mitochondrial network, even with PFA 4% alone, direct perfusion led to a more consistent preservation of mitochondria morphology in vivo (dendritic mitochondria mean length: indirect PFA – 1.95 ± 0.1μm, direct PFA - 4.5 ± 0.4μm, indirect PFA/GA – 1.5 ± 0.1μm, direct PFA/GA - 5.4 ± 0.5μm), and is clearly a critical step in capturing the in vivo structure of mitochondria.

### Anesthesia without oxygen supply during perfusion induces mitochondrial fragmentation

We confirmed that direct perfusion could preserve mitochondrial morphology during perfusion. However, variance in dendritic mitochondrial length is still observed in previous studies, and also in our experiments while using direct perfusion. This suggested that there could be another important factor affecting mitochondrial morphology. Because oxygen levels in mice may change depending on the oxygen supply during anesthesia, and it has been reported that oxygen deprivation can induce mitochondrial fragmentation in several cell types, we examined if anesthesia with and without oxygen supply would result in differences in mitochondrial morphology. In order to study mitochondrial morphology after perfusion, we sparsely labeled mouse cortical neurons using IUE with a Cre-dependent plasmid containing a mitochondria-targeted fluorescent protein and low concentration of a Cre recombinase expressing plasmid (**Figure S2**). At postnatal day (P)21, mice were anesthetized either with an anesthesia machine, which delivers isoflurane with oxygen, or with a closed jar containing isoflurane and directly perfused with 2% PFA/0.075% GA. Oxygen concentration and pulse rate were monitored with a pulse oximeter during perfusion. Anesthesia without oxygen supplement showed significantly decreased oxygen concentration (86.78% ± 1.00% with oxygen vs 55.68% ± 1.93% without oxygen) and increased pulse rate compared to the with oxygen condition (337.1 ± 5.89 pulses/min with oxygen vs 459.5 ± 3.36 pulses/min without oxygen, **Figure 3D-E**). Next, we compared the length of dendritic mitochondria in each condition. Strikingly, mice anesthetized without oxygen showed significantly shortened dendritic mitochondria after perfusion (2.30 ± 0.20μm without oxygen vs 6.74 ± 0.51μm with oxygen, **Figure 3A-C**). These results suggest that avoidance of hypoxic condition is critical for preserving mitochondrial morphology of neurons in vivo.

### Reduced oxygen level causes changes of mitochondrial morphology in vitro

To ascertain what level of hypoxia can result in mitochondrial fragmentation, we next tested if gradual changes in oxygen level could directly affect mitochondrial morphology. Live imaging of fluorescently mitochondria-labeled cortical neurons in vitro allowed us to observe the changes in mitochondrial morphology in real-time. We incubated neurons under normoxic (20% oxygen) and hypoxic conditions (10%, 5%, 0.1% oxygen). As oxygen level dropped, the length of dendritic mitochondria gradually decreased (from 10.21 ± 0.64μm in 20% oxygen to 6.96 ± 0.46μm in 0.1% oxygen, **Figure 4**). Coupled with our previous in vivo perfusion observations, these results emphasize the importance of oxygen concentration for maintaining mitochondrial morphology during fixation.

**Figure 4:**
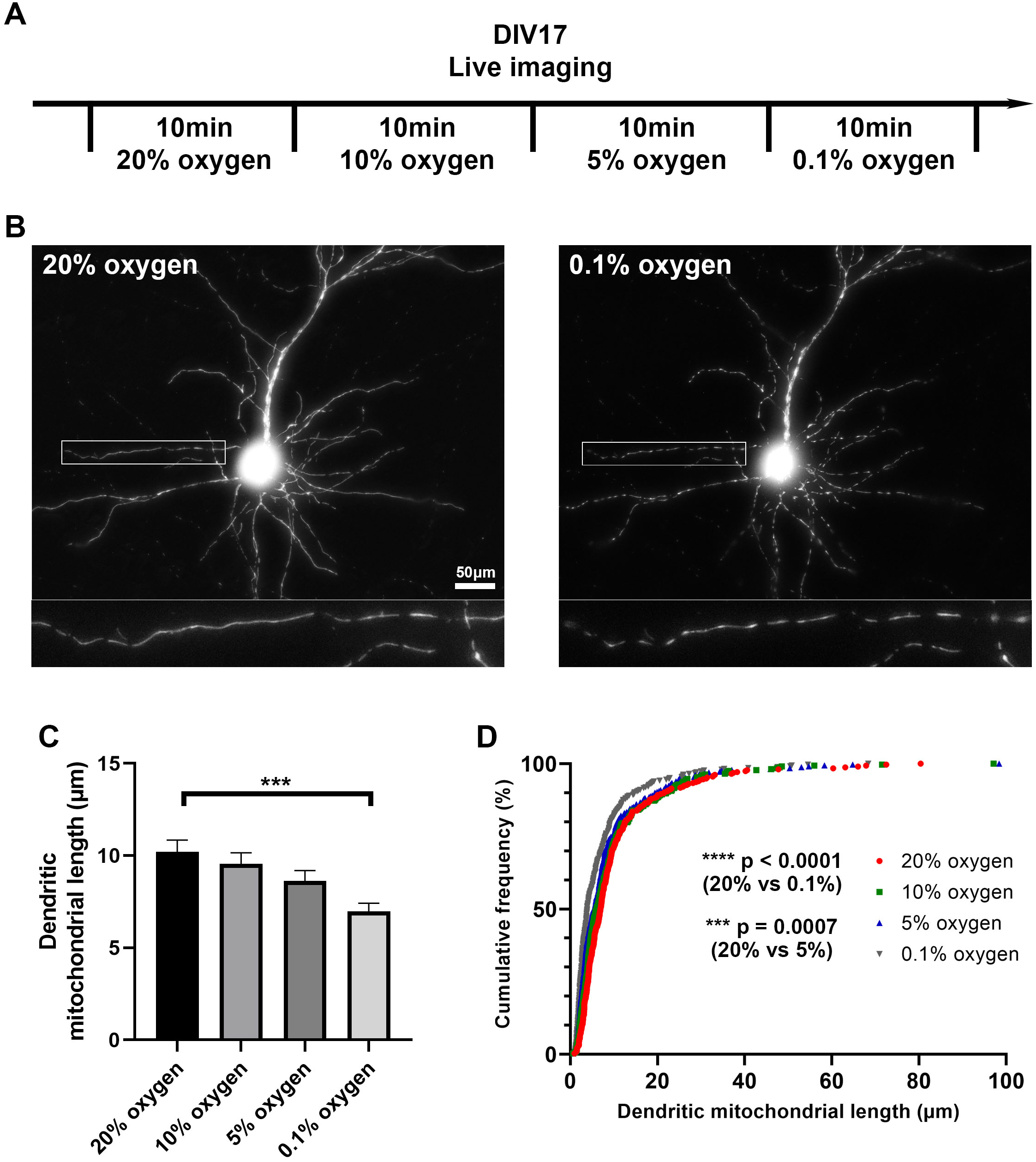
Low oxygen level induces mitochondrial fragmentation in vitro. A) Schematic diagram of live imaging. Live imaging of mitochondria-labeled cortical neurons was done in 4 different oxygen levels; 20%, 10%, 5%, 0.1%. B) Representative images of mitochondria in 20% and 0.1% oxygen level. C) Quantification of dendritic mitochondrial length showed gradual decrease as oxygen level drops. D) Cumulative frequencies of dendritic mitochondrial length also showed shorter mitochondria in low oxygen level, compared to normal oxygen level. n_20% oxygen_ = 2 cultures, 26 dendrites, 316 mitochondria, n_10% oxygen_ = 2 cultures, 26 dendrites, 323 mitochondria, n_5% oxygen_ = 2 cultures, 26 dendrites, 330 mitochondria, no_0.1% oxygen_ = 2 cultures, 26 dendrites, 346 mitochondria. Kruskal–Wallis test with Dunn’s multiple comparisons test. length: *** p=0.0007 for 20% oxygen vs 5% oxygen, **** p<0.0001 for 20% oxygen vs 0.1% oxygen. Scale bar = 20μm

### Fixation protocol affects the outcome of mitochondrial analysis in neurodegenerative disease models

Mitochondrial dysfunction is a hallmark of neurodegeneration. Fragmentation of dendritic mitochondria is observed in many neurodegenerative diseases including Alzheimer’s disease and Parkinson’s diseases (Wang, Su et al. 2009, Matheoud, Sugiura et al. 2016, Zhang, Trushin et al. 2016, Burbulla, Song et al. 2017, Lee, Hirabayashi et al. 2018). As shown in **Figures 1-2**, fixation with 4% PFA resulted in fragmentation of mitochondria, while 2% PFA with 0.075% GA showed intact morphology. Because different compositions of fixative could alter mitochondrial morphology, we assumed that this could influence the analysis of mitochondria in models of neurodegenerative disease. To test this, we investigated the effect of three different fixation solutions using an α-synuclein preformed fibril (PFF)-induced synucleinopathy model. Neurons were incubated with α-synuclein for 11 days starting from 10 DIV and then fixed with 3 different fixatives; 4% PFA, 2% PFA with 0.075% GA, or 4% PFA with 4% sucrose. α-synuclein fibril accumulation was confirmed with phosphorylated α-synuclein staining, which is a marker for accumulated α-synuclein in cells (**Figure S3**). To determine if the fixation solution could affect the analysis outcome of mitochondrial morphology, we compared the differences in dendritic mitochondrial length between control and α-synuclein treated samples in each fixation condition. With 2% PFA/0.075% GA, control mitochondrial morphology was consistent with the results presented above, while α-synuclein treated samples showed a significant decrease in average dendritic mitochondrial length (2.80 ± 0.09μm vs 3.35 ± 0.13μm in control, **Figure 5**). However, fixation with only 4% PFA abolished the differences observed in dendritic mitochondrial length following α-synuclein treatment (2.26 ± 0.07μm vs 2.17 ± 0.06μm in control, **Figure 5**) as control mitochondria were much shorter than with PFA/GA. Although PFA/GA fixation was ideal for preserving mitochondrial morphology, there were high background autofluorescence signals following immunocytochemistry. Thus, we also tested 4% PFA/4% sucrose as an alternative solution, which is known to be cryoprotective. Compared with PFA only, PFA with sucrose preserved mitochondrial morphology closer to PFA/GA fixation but with lower background signals (**Figures 5B, S3**). Overall, these results suggest that fixation solutions could affect the results of mitochondrial morphology analysis, and that this parameter should be carefully weighed during the experimental design phase.

**Figure 5:**
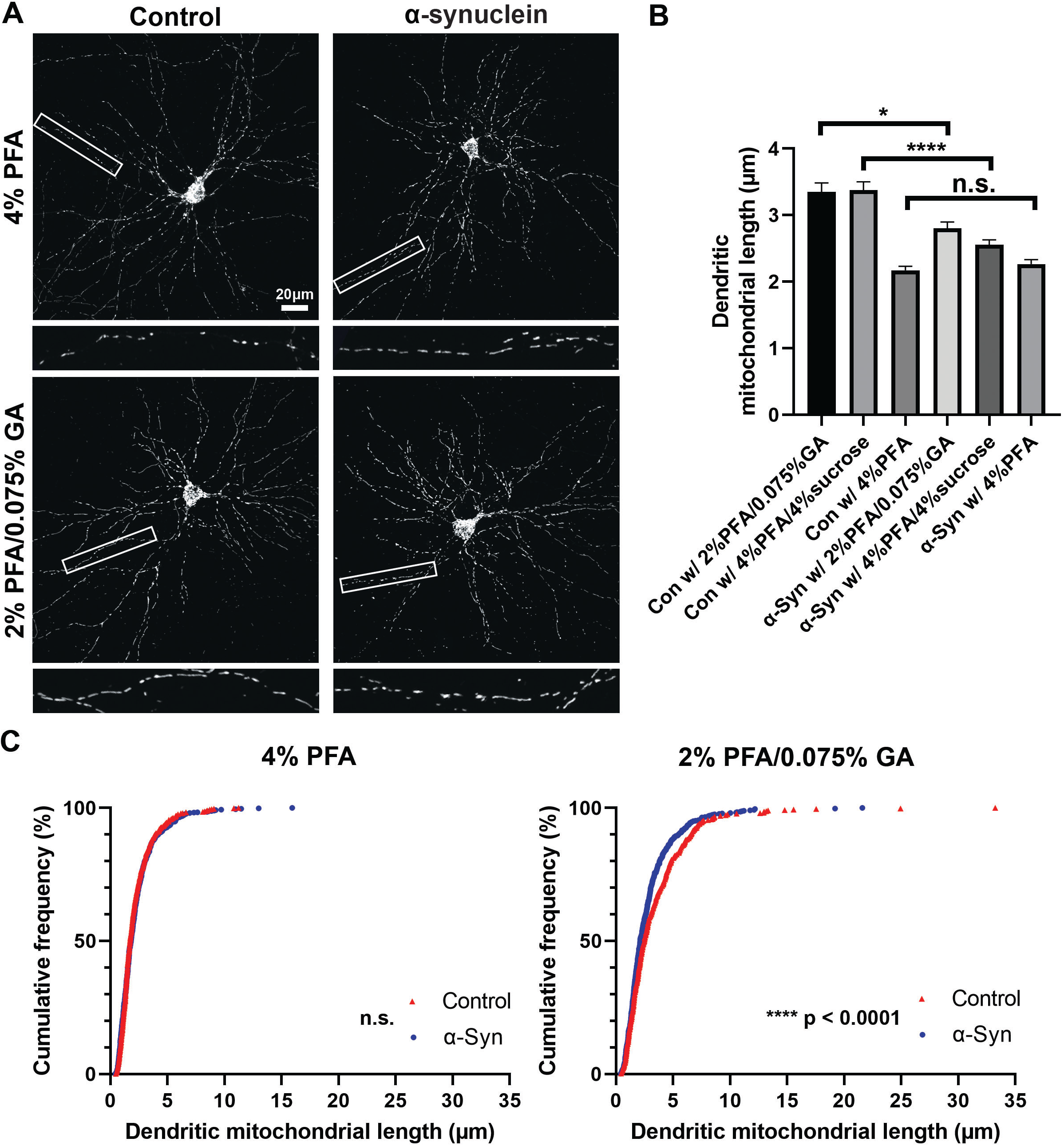
Fixative affects mitochondrial morphology analysis of a neurodegenerative disease model. A) Images of control and α-Syn-treated neurons, which were fixed with three different fixation solutions (4% PFA, 2% PFA/0.075% GA, 4% PFA/4% sucrose) for 15 min at DIV19 B) Quantification of dendritic mitochondrial length confirmed 2% PFA/0.075% GA and 4% PFA/4% sucrose fixed samples showed differences between α-Syn-treated neurons and control neurons. In contrast, there were no differences in 4% PFA fixed samples. n_Con w/ 4%PFA_ = 3 cultures, 34 dendrites, 510 mitochondria, n_Con w/ PFA/GA_ = 3 cultures, 32 dendrites, 442 mitochondria, n_Con w/ PFA/Sucrose_ = 3 cultures, 32 dendrites, 625 mitochondria, n_α-syn w/ 4%PFA_ = 3 cultures, 32 dendrites, 587 mitochondria, n_α-syn w/ PFA/GA_ = 3 cultures, 34 dendrites, 642 mitochondria, n_α-syn w/ PFA/Sucrose_ = 3 cultures, 36 dendrites, 629 mitochondria. Kruskal-Wallis test with Dunn’s multiple comparisons test, * p=0.03 for control vs α-Syn in 2% PFA/0.075% GA, **** p<0.0001 for control vs α-Syn in 4% PFA/4% Sucrose. C) Cumulative frequencies of dendritic mitochondrial length in 4% PFA showing no differences in dendritic mitochondria. D) Cumulative frequencies of dendritic mitochondrial length in 2% PFA/0.075% GA fixation.

## DISCUSSION

Our results clearly demonstrate that the method of fixation is critical for preserving the mitochondria morphology observed in living mammalian neurons. Our findings can be broken down into three main observations: (1) direct fixation is critical both in cultured neurons and in vivo. Any condition which included significant pre-washing with PBS or other saline solutions to remove culture medium or blood resulted in fragmentation of the mitochondria. (2) preventing hypoxia is required to maintain mitochondrial structure. Conditions that resulted in low oxygen were sufficient to induce mitochondrial fragmentation both in vitro and in vivo. (3) a mixture of PFA/GA most faithfully preserved the mitochondrial morphologies and sizes observed in living neurons. This appears especially true with cultured neurons as even direct fixation with 4% PFA resulted in fragmented mitochondria. However in vivo, if conditions one and two are met we observed only a small decrease in mitochondrial length with 4% PFA compared to 2% PFA/0.075% GA. Together, our results establish that it is critical for the fixation process to occur as rapidly as possible in order to maintain the mitochondria structure observed in living neurons, and if this is not carefully considered the fixative conditions may result in incorrect conclusions as we have shown with α-synuclein treatment.

Surprisingly, “standard” published protocols for the perfusion of rodents and other small animals show considerable variability (Buffalo, Montana, Gage, Kipke et al. 2012, Uhlig, Krause et al. 2015, Wu, Cai et al. 2021, Chu 2022). This variability mainly comes in two forms: the anesthesia used to prepare the animal for the procedure, and the buffers and/or fixatives used during the procedure. It is clear that the choices made at each of these steps will have important ramifications on the outcome of the procedure, and the decision should depend on the ultimate focus of the investigator. For instance, in our hands overall neuronal cell structure (i.e. dendrites, axons, spines) appeared to be well preserved with all the anesthetics and fixatives we tested, even though mitochondrial structure showed the striking differences detailed above (data not shown).

While future work is still required to fully understand the mechanism, we found that exposing mitochondria to a short-term hypoxic environment is sufficient to induce mitochondrial fragmentation. Molecularly this could be through mTOR-Drp1 mediated activation or by increased FUNDC1-Drp1 interaction (Wu, Lin et al. 2016, Zheng, Qian et al. 2019). Interestingly, in vivo hypoxic conditions triggered more significant mitochondrial fragmentation (less than 2 min) than in cultured neurons, which might be caused by the fast delivery of low oxygen blood and/or solutions. Both the mode of anesthesia and the mode of perfusion likely play a role in the induction of this hypoxic state. Our results argue that when studying mitochondria, methods of anesthesia resulting in low oxygen levels should be avoided (i.e. CO2, drop isofluorane). Recent work demonstrates that oxygen supplementation is required to prevent hypoxia with both inhaled and injected anesthetics (Blevins, Celeste et al. 2021). One step that is preserved across the majority of published perfusion protocols is a saline based flush (Buffalo, Montana, Gage, Kipke et al. 2012, Wu, Cai et al. 2021, Chu 2022). While this step removes blood, it likely exacerbates the hypoxia in the sample and should clearly be avoided when downstream analysis of mitochondria morphology will be performed.

Two recent papers (Qin, Jiang et al. 2021, Hinton, Katti et al. 2022) have independently come to some of the same conclusions about the role of fixation on mitochondria structure; nonetheless, key differences exist between in our findings. Hinton et al used electron microscopy to show that anesthesia with an isoflurane/oxygen mixture provided better preservation of mitochondrial structure than CO_2_-based methods. In addition, direct perfusion of 4% PFA compared to PBS flushing conditions after injectable anesthesia (ketamine/xylazine) maintained mitochondrial shape. However, both the oxygen supplement and direct perfusion method were followed by strong fixative (Trump’s solution; 4% formaldehyde/1% GA) immersion, and actual oxygen level was not monitored. Our study combined the oxygen supplementation and direct perfusion of fixative, and also checked and manipulated oxygen level in vivo and in vitro. Therefore, we could unambiguously conclude that the hypoxic condition caused the fragmentation of mitochondria. While Qin et al show that PFA/GA mixtures preserve mitochondria structure better via fluorescence imaging, much higher concentrations of GA were tested (2.5%-1.5%) and the experiments were performed in MEFs not neurons. High concentrations of GA cause strong background signals during immunocytochemistry, thus we find that the combination of low GA concentration coupled with the oxygenated/direct perfusion allows for both the preservation of mitochondria structure and for immunocytochemistry in the same brain samples.

Our results also argue that we should carefully consider the role that the method of fixation may have played in findings about mitochondrial morphology or structure. This may impact the outcome in a few different ways. One situation would be an incorrect negative result where incorrect perfusion leads to all conditions giving rise to similarly highly fragmented mitochondria as a result of a slow fixation process (**Figure 5**). In another situation, sub-optimal fixation conditions could give rise to a false positive result. For instance, under a pathogenic situation, stressed mitochondria could be structurally similar in the living cells but are more sensitive to the fixation process leading to fixation induced fragmentation only in the stressed mitochondria. Clearly the gold standard would be to visualize mitochondrial structure under living conditions which would remove the potential for fixation artifacts.

A potential confounding factor in the variation observed in the literature regarding mitochondrial length is recent data reporting differential mitochondria morphologies in different brain regions. It is now clear that both distinct neuron types and even the compartments within the same neuron (soma, dendrites, axon) regulate mitochondria morphology to different levels (Faits, Zhang et al. 2016, Lewis, Kwon et al. 2018, Chandra, Calarco et al. 2019, Janickova, Rechberger et al. 2020, Faitg, Lacefield et al. 2021, Lee, Kondapalli et al. 2022). However, it is unclear if or how these different mitochondrial populations would be affected by the different fixatives and perfusion methods.

In each condition that we tested, a fixative solution of PFA/GA maintained mitochondrial morphology closer to those observed in living neurons. However, fixing with PFA/GA does have potential drawbacks that may limit its usefulness based on the context of the experiment. First, PFA/GA fixation causes increased background fluorescence across the visible wavelengths. Therefore, if trying to visualize a lowly expressed protein the high background may reduce the signal to noise ratio to a level that is unacceptable. Finally for antibodies that still work after PFA/GA fixation, incubation times need to be significantly increased presumably because of the increased crosslinking that occurs with the PFA/GA fixation. We routinely had to double our antibody incubation times for 100μm thick brain sections from 16-24hrs with PFA only to 32-48hrs following PFA/GA fixation. In addition, the use of a 4% PFA/4% sucrose fixative solution showed less background signal without affecting dendritic mitochondrial length in vitro (**Figures 5, S3**); therefore, this may also be considered for immunocytochemistry experiments.

Taken together, our results provide standard anesthesia and fixation methods for the study of neuronal mitochondrial morphology both in vitro and in vivo. These results can also serve as a starting point when investigating other cellular mechanisms that are vulnerable to environmental stress.

## Abbreviations

PN: pyramidal neurons
PFA: paraformaldehyde
GA: glutaraldehyde
ATP: adenosine triphosphate
EM: electron microscopy
EUE: ex utero electroporation
IUE: in utero electroporation

## Acknowledgements

We thank Drs. Julien Courchet, Yusuke Hirabayashi and Franck Polleux for discussions and critical feedback on the manuscript. We also thank all members of the Kwon and Lewis labs for feedback and discussion along the way. This research was supported by the National Research Foundation (NRF) funded by the Korean government (MSIT) (2019M3E5D2A01063794, 2020R1C1C1006386, 2022M3E5E8017395), and KIST Program (2E31511) to SK Kwon, and OMRF startup funds and support from the Presbyterian Health Foundation to TL Lewis.

## Author Contributions

Conceptualization: SKK & TL. Formal analysis: SYK, KS, BO, SKK, TL. Funding acquisition: SKK, TL. Investigation: SYK, KS, BO, SKK, TL. Methodology: SYK, KS, BO, SKK, TL. Project administration: KH, SKK, TL. Resources: SKK, TL. Supervision: KH, SKK, TL. Visualization: SYK, KS, BO, SKK, TL. Writing – original draft: SYK, SKK, TL. Writing – review & editing: SYK, KS, BO, KH, SKK, TL.

---

**Figure S-1:**
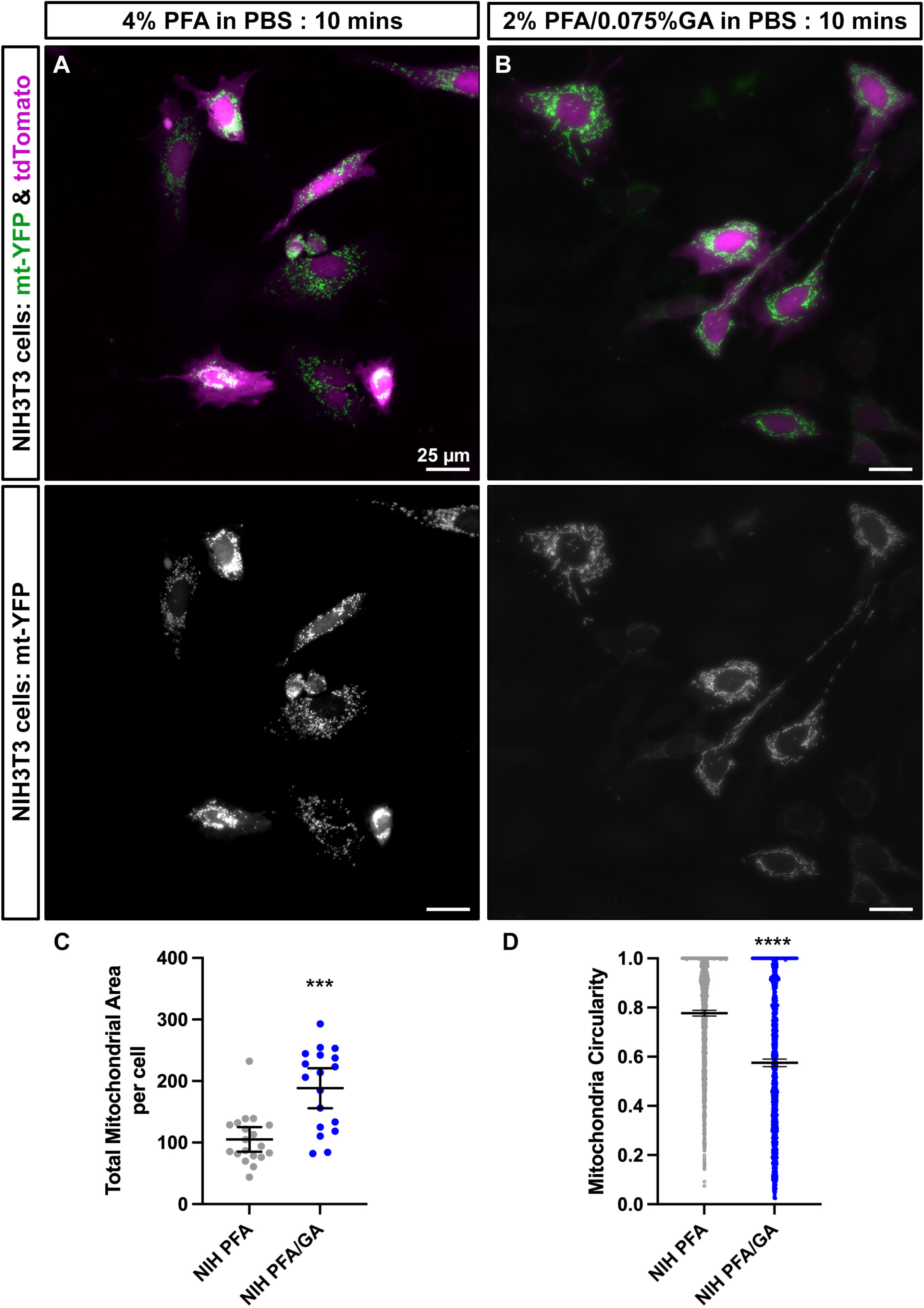
combination of paraformaldehyde and glutaraldehyde maintains mitochondria morphology better than paraformaldehyde alone in cultured NIH3T3 cells. A) Representative NIH3T3 cells labeled with mitochondrial matrix targeted YFP (mt-YFP, green) and tdTomato (purple) imaged after fixation with 4% PFA in PBS. B) Representative NIH3T3 cells labeled with mt-YFP (green) and tdTomato (purple) imaged after fixation with 2% PFA/0.075% GA in PBS. C) Quantification of total mitochondrial area per cell showing that fixation with PFA along causes mitochondrial fragmentation (mean +/− 95% CI). D) Quantification of individual mitochondria for circularity shown as mean +/− 95% CI before and after fixation with the indicated fixative solution. Mann-Whitney test. n_4%PFAcells_= 19 cells, n_PFA/GAcells_= 18 cells, n_4%PFAcircularity_= 1347 mitochondria, n_PFA/GAcircularity_= 1400 mitochondria. p*** p<0.001, **** p<0.0001. Scale bar = 25μm

**Figure S2:**
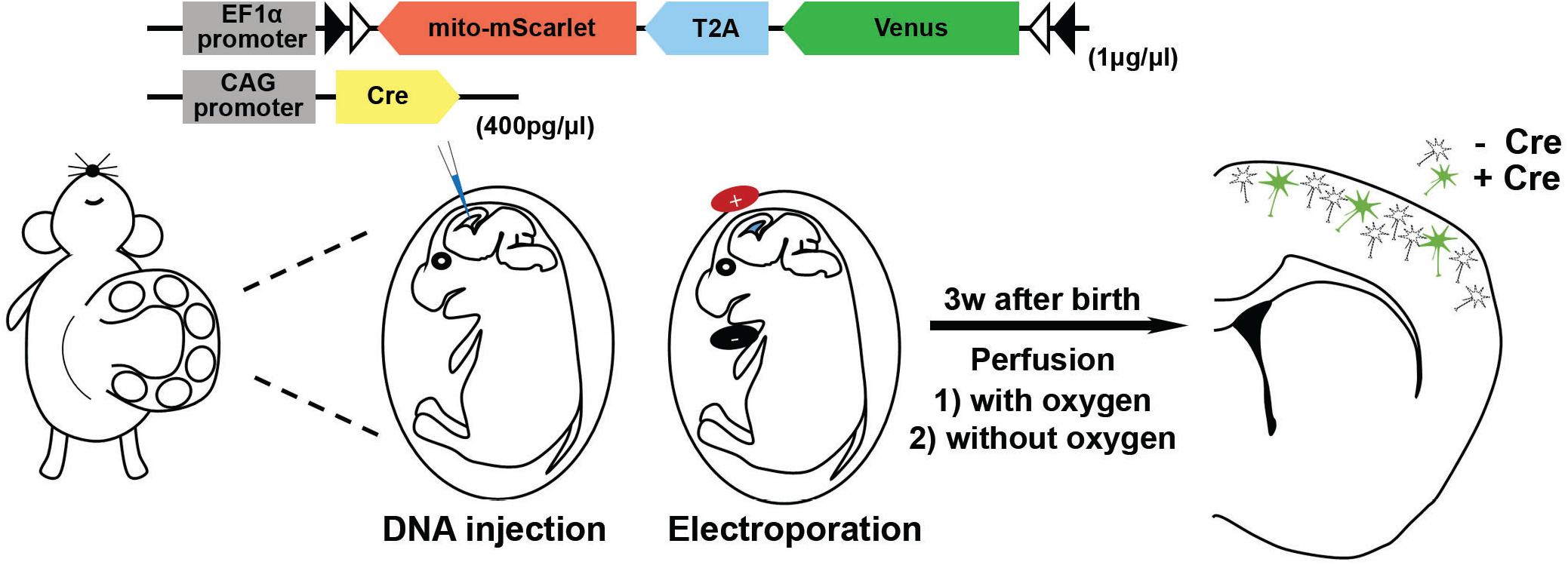
In utero electroporation strategy for sparse labeling of cortical neurons in vivo. A “DIO” plasmid encoding mitochondrial targeted mScarlet and cytoplasmic Venus was mixed with a low concentration of a second plasmid encoding Cre and then electroporated in E15.5 pups to give sparse labeling of cortical neurons in vivo.

**Figure S3:**
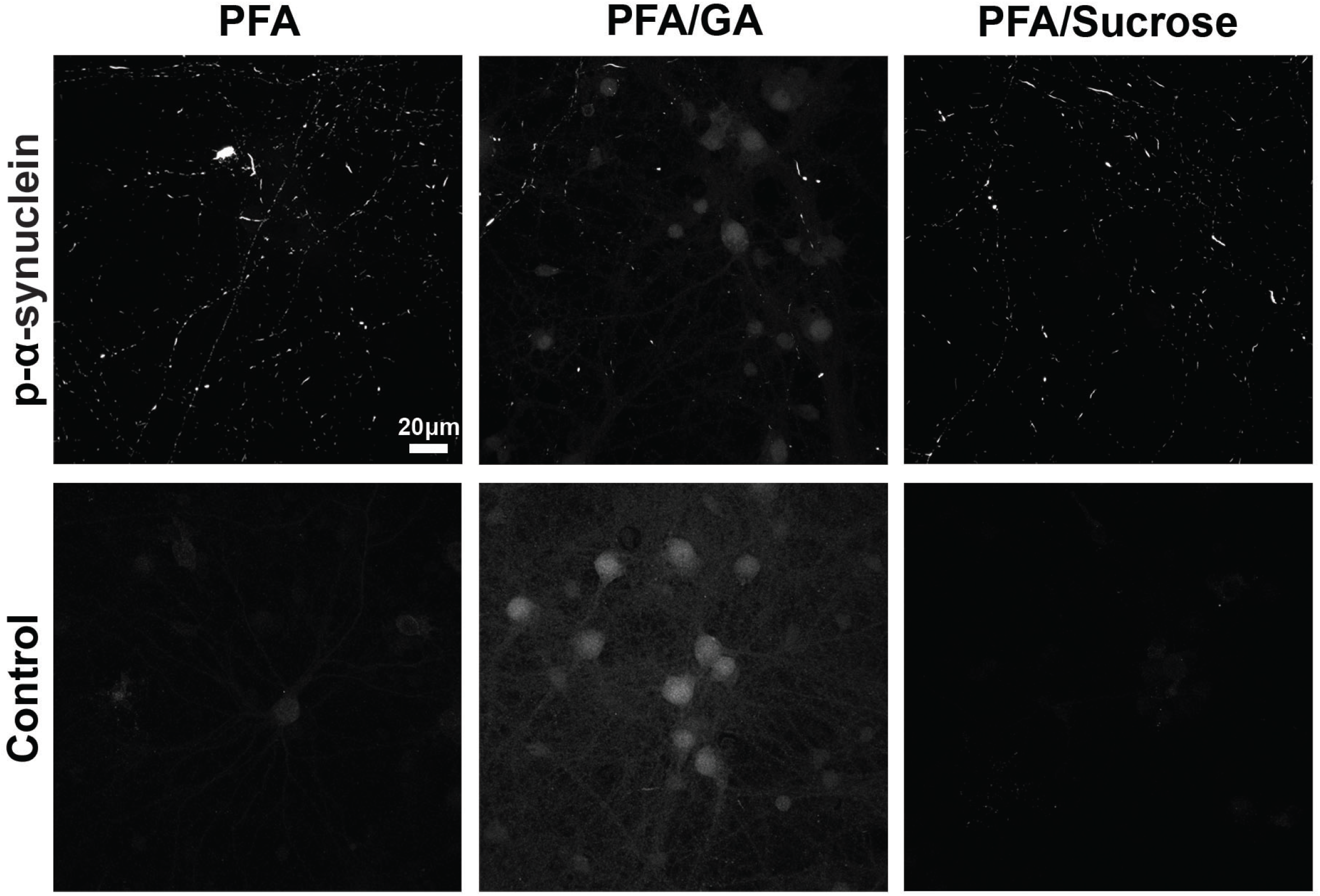
p-α-synuclein staining in different fixation solutions. α-Syn accumulation was confirmed with p-α-Syn staining. Both in control and α-Syn treated samples, 2% PFA/0.075% GA-fixed images showed higher background signals compared to 4% PFA or 4% PFA/4% sucrose fixatives. Scale bar = 20μm

## References

Azzu, V. and T. G. Valencak (2017). “Energy Metabolism and Ageing in the Mouse: A Mini-Review.” Gerontology 63(4): 327–336.

Belanger, M., I. Allaman and P. J. Magistretti (2011). “Brain energy metabolism: focus on astrocyte-neuron metabolic cooperation.” Cell Metab 14(6): 724–738.

Blevins, C. E., N. A. Celeste and J. O. Marx (2021). “Effects of Oxygen Supplementation on Injectable and Inhalant Anesthesia in C57BL/6 Mice.” J Am Assoc Lab Anim Sci 60(3): 289–297.

Buffalo, U. o. “Standard Operating Procedures For Whole Body Perfsuion Fixation Of Mice.” Retrieved 09/02/22, from https://www.buffalo.edu/content/dam/www/research/pdf/laf/sop/2A12.pdf.

Burbulla, L. F., P. Song, J. R. Mazzulli, E. Zampese, Y. C. Wong, S. Jeon, D. P. Santos, J. Blanz, C. D. Obermaier, C. Strojny, J. N. Savas, E. Kiskinis, X. Zhuang, R. Kruger, D. J. Surmeier and D. Krainc (2017). “Dopamine oxidation mediates mitochondrial and lysosomal dysfunction in Parkinson’s disease.” Science 357(6357): 1255–1261.

Chandel, N. S. (2014). “Mitochondria as signaling organelles.” BMC Biol 12: 34.

Chandra, R., C. A. Calarco and M. K. Lobo (2019). “Differential mitochondrial morphology in ventral striatal projection neuron subtypes.” J Neurosci Res 97(12): 1579–1589.

Chang, D. T. W. and I. J. Reynolds (2006). “Differences in mitochondrial movement and morphology in young and mature primary cortical neurons in culture.” Neuroscience 141(2): 727–736.

Chu, H. Y. (2022). “Standard Operating Procedure: Mouse transcardiac perfusion protocol.” protocols.io.

Diaz-Garcia, C. M. and G. Yellen (2019). “Neurons rely on glucose rather than astrocytic lactate during stimulation.” J Neurosci Res 97(8): 883–889.

Faitg, J., C. Lacefield, T. Davey, K. White, R. Laws, S. Kosmidis, A. K. Reeve, E. R. Kandel, A. E. Vincent and M. Picard (2021). “3D neuronal mitochondrial morphology in axons, dendrites, and somata of the aging mouse hippocampus.” Cell Rep 36(6): 109509.

Faits, M. C., C. Zhang, F. Soto and D. Kerschensteiner (2016). “Dendritic mitochondria reach stable positions during circuit development.” Elife 5: e11583.

Fox, P. T., M. E. Raichle, M. A. Mintun and C. Dence (1988). “Nonoxidative glucose consumption during focal physiologic neural activity.” Science 241(4864): 462–464.

Gage, G. J., D. R. Kipke and W. Shain (2012). “Whole animal perfusion fixation for rodents.” J Vis Exp (65).

Gonzalez-Rodriguez, P., E. Zampese, K. A. Stout, J. N. Guzman, E. Ilijic, B. Yang, T. Tkatch, M. A. Stavarache, D. L. Wokosin, L. Gao, M. G. Kaplitt, J. Lopez-Barneo, P. T. Schumacker and D. J. Surmeier (2021). “Disruption of mitochondrial complex I induces progressive parkinsonism.” Nature 599(7886): 650–656.

Hinton, A., P. Katti, T. A. Christensen, M. Mungai, J. Shao, L. Zhang, S. Trushin, A. Alghanem, A. Jaspersen, R. E. Geroux, K. Neikirk, M. Biete, E. G. Lopez, Z. Vue, H. K. Beasley, A. G. Marshall, J. Ponce, C. K. E. Bleck, I. Hicsasmaz, S. A. Murray, R. A. C. Edmonds, A. Dajles, Y. D. Koo, S. Bacevac, J. L. Salisbury, R. O. Pereira, B. Glancy, E. Trushina and E. D. Abel (2022). “A comprehensive approach to artifact-free sample preparation and the assessment of mitochondrial morphology in tissue and cultured cells.” bioRxiv: 2021.2006.2027.450055.

Janickova, L., K. F. Rechberger, L. Wey and B. Schwaller (2020). “Absence of parvalbumin increases mitochondria volume and branching of dendrites in inhibitory Pvalb neurons in vivo: a point of convergence of autism spectrum disorder (ASD) risk gene phenotypes.” Mol Autism 11(1): 47.

Kimura, T. and F. Murakami (2014). “Evidence that dendritic mitochondria negatively regulate dendritic branching in pyramidal neurons in the neocortex.” J Neurosci 34(20): 6938–6951.

Kwon, S. K., R. Sando, 3rd, T. L. Lewis, Y. Hirabayashi, A. Maximov and F. Polleux (2016). “LKB1 Regulates Mitochondria-Dependent Presynaptic Calcium Clearance and Neurotransmitter Release Properties at Excitatory Synapses along Cortical Axons.” PLoS Biol 14(7): e1002516.

Lee, A., Y. Hirabayashi, S. K. Kwon, T. L. Lewis, Jr. and F. Polleux (2018). “Emerging roles of mitochondria in synaptic transmission and neurodegeneration.” Curr Opin Physiol 3: 82–93.

Lee, A., C. Kondapalli, D. M. Virga, T. L. Lewis, Jr., S. Y. Koo, A. Ashok, G. Mairet-Coello, S. Herzig, M. Foretz, B. Viollet, R. Shaw, A. Sproul and F. Polleux (2022). “Abeta42 oligomers trigger synaptic loss through CAMKK2-AMPK-dependent effectors coordinating mitochondrial fission and mitophagy.” Nat Commun 13(1): 4444.

Lewis, T. L., Jr., S. K. Kwon, A. Lee, R. Shaw and F. Polleux (2018). “MFF-dependent mitochondrial fission regulates presynaptic release and axon branching by limiting axonal mitochondria size.” Nat Commun 9(1): 5008.

Lewis, T. L., Jr., G. F. Turi, S. K. Kwon, A. Losonczy and F. Polleux (2016). “Progressive Decrease of Mitochondrial Motility during Maturation of Cortical Axons In Vitro and In Vivo.” Curr Biol 26(19): 2602–2608.

Lin, Y., L. L. Li, W. Nie, X. Liu, A. Adler, C. Xiao, F. Lu, L. Wang, H. Han, X. Wang, W. B. Gan and H. Cheng (2019). “Brain activity regulates loose coupling between mitochondrial and cytosolic Ca(2+) transients.” Nat Commun 10(1): 5277.

Matheoud, D., A. Sugiura, A. Bellemare-Pelletier, A. Laplante, C. Rondeau, M. Chemali, A. Fazel, J. J. Bergeron, L. E. Trudeau, Y. Burelle, E. Gagnon, H. M. McBride and M. Desjardins (2016). “Parkinson’s Disease-Related Proteins PINK1 and Parkin Repress Mitochondrial Antigen Presentation.” Cell 166(2): 314–327.

Montana, U. o. “Rodent Brain Perfusion.” Retrieved 09/02/2022, from https://www.umt.edu/research/LAR/sops/sop-perfusion.php.

Owens, K., J. H. Park, S. Gourley, H. Jones and T. Kristian (2015). “Mitochondrial dynamics: cell-type and hippocampal region specific changes following global cerebral ischemia.” J Bioenerg Biomembr 47(1-2): 13–31.

Popov, V., N. I. Medvedev, H. A. Davies and M. G. Stewart (2005). “Mitochondria form a filamentous reticular network in hippocampal dendrites but are present as discrete bodies in axons: a three-dimensional ultrastructural study.” J Comp Neurol 492(1): 50–65.

Qin, Y., W. Jiang, A. Li, M. Gao, H. Liu, Y. Gao, X. Tian and G. Gong (2021). “The Combination of Paraformaldehyde and Glutaraldehyde Is a Potential Fixative for Mitochondria.” Biomolecules 11(5).

Rangaraju, V., M. Lauterbach and E. M. Schuman (2019). “Spatially Stable Mitochondrial Compartments Fuel Local Translation during Plasticity.” Cell 176(1-2): 73–84 e15.

Rolfe, D. F. and G. C. Brown (1997). “Cellular energy utilization and molecular origin of standard metabolic rate in mammals.” Physiol Rev 77(3): 731–758.

Swerdlow, R. H., J. M. Burns and S. M. Khan (2014). “The Alzheimer’s disease mitochondrial cascade hypothesis: progress and perspectives.” Biochim Biophys Acta 1842(8): 1219–1231.

Turner, N. L., T. Macrina, J. A. Bae, R. Yang, A. M. Wilson, C. Schneider-Mizell, K. Lee, R. Lu, J. Wu, A. L. Bodor, A. A. Bleckert, D. Brittain, E. Froudarakis, S. Dorkenwald, F. Collman, N. Kemnitz, D. Ih, W. M. Silversmith, J. Zung, A. Zlateski, I. Tartavull, S. C. Yu, S. Popovych, S. Mu, W. Wong, C. S. Jordan, M. Castro, J. Buchanan, D. J. Bumbarger, M. Takeno, R. Torres, G. Mahalingam, L. Elabbady, Y. Li, E. Cobos, P. Zhou, S. Suckow, L. Becker, L. Paninski, F. Polleux, J. Reimer, A. S. Tolias, R. C. Reid, N. M. da Costa and H. S. Seung (2022). “Reconstruction of neocortex: Organelles, compartments, cells, circuits, and activity.” Cell 185(6): 1082–1100 e1024.

Uhlig, C., H. Krause, T. Koch, M. Gama de Abreu and P. M. Spieth (2015). “Anesthesia and Monitoring in Small Laboratory Mammals Used in Anesthesiology, Respiratory and Critical Care Research: A Systematic Review on the Current Reporting in Top-10 Impact Factor Ranked Journals.” PLoS One 10(8): e0134205.

Wang, X., B. Su, H. G. Lee, X. Li, G. Perry, M. A. Smith and X. Zhu (2009). “Impaired balance of mitochondrial fission and fusion in Alzheimer’s disease.” J Neurosci 29(28): 9090–9103.

Wu, J., Y. Cai, X. Wu, Y. Ying, Y. Tai and M. He (2021). “Transcardiac Perfusion of the Mouse for Brain Tissue Dissection and Fixation.” Bio Protoc 11(5): e3988.

Wu, W., C. Lin, K. Wu, L. Jiang, X. Wang, W. Li, H. Zhuang, X. Zhang, H. Chen, S. Li, Y. Yang, Y. Lu, J. Wang, R. Zhu, L. Zhang, S. Sui, N. Tan, B. Zhao, J. Zhang, L. Li and D. Feng (2016). “FUNDC1 regulates mitochondrial dynamics at the ER-mitochondrial contact site under hypoxic conditions.” EMBO J 35(13): 1368–1384.

Zhang, L., S. Trushin, T. A. Christensen, B. V. Bachmeier, B. Gateno, A. Schroeder, J. Yao, K. Itoh, H. Sesaki, W. W. Poon, K. H. Gylys, E. R. Patterson, J. E. Parisi, R. Diaz Brinton, J. L. Salisbury and E. Trushina (2016). “Altered brain energetics induces mitochondrial fission arrest in Alzheimer’s Disease.” Sci Rep 6: 18725.

Zheng, X., Y. Qian, B. Fu, D. Jiao, Y. Jiang, P. Chen, Y. Shen, H. Zhang, R. Sun, Z. Tian and H. Wei (2019). “Mitochondrial fragmentation limits NK cell-based tumor immunosurveillance.” Nat Immunol 20(12): 1656–1667.

Zhou, X., H. Chen, L. Wang, C. Lenahan, L. Lian, Y. Ou and Y. He (2021). “Mitochondrial Dynamics: A Potential Therapeutic Target for Ischemic Stroke.” Front Aging Neurosci 13: 721428.

